# Bioinformatics analysis of endophytic bacteria related to berberine in the Chinese medicinal plant *Coptis teeta* Wall

**DOI:** 10.1101/760777

**Authors:** Tian-hao Liu, Xiao-mei Zhang, Shou-zheng Tian, Li-guo Chen, Jia-li Yuan

**Author notes:** Corresponding author: Shou-zheng Tian; No. 1076 Yuhua Road, Chenggong District 650500, Kunming, Yunnan, China.

## Abstract

Plant endophytic microorganisms absorb nutrients and prevent pathogen damage, supporting healthy plant growth. However, relationships between endophytic bacteria of the medicinal plant *Coptis teeta* Wall. and berberine production remain unclear. Herein, we explored the microbial composition of wild-type (WT) and cultivated *Coptis teeta* Wall. root, stem and leaf, and endophytic bacteria related to berberine. Microbial characteristics of were analyzed by 16S rDNA sequencing, and berberine in roots was analyzed by high-performance liquid chromatography (HPLC). Proteobacteria, Actinobacteria and Bacteroidetes were the major phyla, and *Mycobacterium, Salmonella, Nocardioides, Burkholderia-Paraburkholderia* and *Rhizobium* were the dominant genera. Berberine was positively correlated with total P (TP), total N (TN), total K (TK) and available K (AK) in rhizosphere soil, and with *Microbacterium* and *norank_f_7B-8*, whereas TK was positively correlated with *Microbacterium*, TN, AK and *Burkholderia-Paraburkholderia*. The findings will support further studies on endophytic bacteria and berberine in *Coptis teeta* Wall., and may promote berberine production.

## Introduction

*Coptis teeta* Wall. belongs to the genus *Ranunculaceae* and is considered an important medicinal plant in the Gaoligongshan area of northwestern Yunnan. In Chinese herbal medicine, *Coptidis rhizome* refers to the dried rhizome of *Coptis chinensis* Franch., *C. deltoidea* CYCheng et Hsiao, and *C. teeta* Wall. [1]. *Coptidis rhizome* is used to treat heat and dampness, detoxification[2], vomiting, diarrhoea, jaundice, high fever, fainting, heartburn, upset stomach, heart palpitations, bloody vomiting, red eyes, toothache, thirst, phlegm, swollen acne, eczema, and various other ailments [3,4]. As long ago as the Ming Dynasty, Lan Mao recorded in the Diannan Materia Medica that *Coptidis rhizome* grown in Yunnan, namely *Coptis teeta* Wall., was more efficacious than that grown in Sichuan. Indeed, *Coptis teeta* Wall. is widely considered to produce the best quality *Coptidis rhizome*.

In addition to use in Chinese medicine, *Coptidis rhizome* is also a source of compounds used in proprietary Chinese pharmaceutical products such as Huanglian 9 and Yinglian tablet. Although statistics are incomplete, there are numerous pharmaceutical preparations containing *Coptidis rhizome* as a raw ingredient. Botanical and pharmacological studies have shown that *Coptidis rhizome* plants contain a variety of alkaloids such as palmatine and the more famous berberine. Studies have shown that the root of *Coptis teeta* Wall. contains 6–7% berberine, while the stem contains 2–3% and the leaf contains 1–1.97%.

*Coptis teeta* Wall. is a shade-tolerant plant that grows at high altitude, in cold mountainous and rainy regions. *Coptis teeta* Wall. is slow-growing, and it takes 6 to 7 years after sowing before products can be harvested. The slow growth is often affected by a variety of diseases and pests, such as anthracnose [5]. Its rhizomes are small, and it produces unusual ‘breeding branches’ that consume a lot of nutrients [6]. Therefore, the yield of *Coptis teeta* Wall. is very low. Furthermore, due to long-term over-harvesting and continuous habitat destruction, the wild population is on the verge of extinction, and the species listed as a second-class endangered plant in China. The growth of *Coptis teeta* Wall. is dependent on various environmental factors including fertiliser, moisture, light and temperature that must be considered when attempting to standardise planting and management to maximise the yield and quality of medicinal materials [7].

Understanding the effects of fertiliser application to rhizosphere soil of medicinal plants is of great significance for improving the yield and ensuring the quality of medicinal materials [8]. Phosphate and compound fertilisers can influence the growth index, yield, and quality of *Coptis teeta* Wall. [9], and N, P and K play important roles in vegetative growth, indicating that their combined application may be key to achieving a high yield. Application of fertiliser can significantly increase the yield, and artificial cultivation of *Coptis teeta* Wall. is the only method capable of meeting demand, hence the need for an improved, standardised approach.

In recent years, plant microbiome research has developed rapidly [10,11]. Interleaf, interstem, rhizosphere and soil microorganisms form a huge planet-wide ecosystem that plays a crucial role in many processes on a global scale. With the development of molecular biology technology, it is possible to study interactions between microbial communities and medicinal components in different parts of medicinal plants. Microorganisms are closely related to plant health; some microorganisms (*Bacillus, Pseudomonas* and others) can reduce the incidence of scab, studies on the relationships between the soil environment and medicinal components of endophytic bacteria in medicinal plants are scarce [12,13].

The present work expands on recent research on the medicinal plant *Coptis teeta* Wall. by investigating the microbial composition in the root, stem and leaf of wild-type (WT) and cultivated *Coptis chinensis*. The synthesis of berberine in *C. chinensis* is likely related to endophytic bacteria, and this is correlated with nutrition. Therefore, in the present study, WT and cultivated *Coptis teeta* Wall. samples were collected from the country of origin, and the microbial characteristics of root, stem and leaf tissues were analysed by 16S rDNA sequencing. Rhizosphere soil samples of were also collected to measure the content of total P (TP), total N (TN), total K (TK) and available K (AK) in the soil. In addition, the berberine content in roots were analysed by high-performance liquid chromatography (HPLC).

## Materials and Methods

### Sample collection and processing

A total of 10 *Coptis teeta* Wall. plants (five WT and five artificially cultivated, referred to as W and A, respectively) were collected from Tengchong (longitude = 98.49, latitude = 25.03). The surface of samples was cleaned with sterile water, disinfected, washed again with sterile water, and different tissues were separated (R = root, S = stem, L = leaf). Corresponding soil samples were collected at the same time as plants and thoroughly air-dried before testing. Samples were numbered 1–5.

### 16s rDNA gene sequencing of endophytic bacteria in roots, stems and leaves

#### DNA extraction and PCR amplification

Extracted genomic DNA was analysed by 1% agarose gel electrophoresis. Specific primers with barcodes were synthesised according to the designated sequencing region and PCR was performed using a TransGen AP221-02 instrument with TransStart Fastpfu DNA Polymerase using standard manufacturer’s procedures. All samples were analysed in triplicate. PCR products were mixed, analysed by 2% agarose gel electrophoresis, and condensed using AxyPrepDNA. A gel recovery kit (AXYGEN) was used to purify excised PCR products. Based on the preliminary quantitative results of electrophoresis, PCR products were quantified using a QuantiFluor-ST Blue Fluorescence Quantification System (Promega), then mixed according to the quantities required for sequencing [14].

#### Miseq library construction and Miseq sequencing

The Illumina official linker sequence was added to the outer end of the target region by PCR, and PCR products were recovered using a gel recovery kit, eluted with TRIS-HCl buffer, analysed by 2% agarose electrophoresis, and hydrolysed with sodium hydroxide. A single-stranded DNA fragment was produced using a TruSeq DNA Sample Prep Kit.

One end of the DNA fragment was complementary to the primer base and immobilised on the chip. Using the DNA fragment as a template, the base sequence immobilised on the chip served as a primer for PCR synthesis, and the target DNA fragment to be detected was synthesised on the chip. After denaturation and annealing, the other end of the DNA fragment on the chip was randomly complementary to another primer in the vicinity, and also fixed to form a ‘bridge’. PCR amplification produces DNA clusters, and the DNA amplicon was linearised into a single strand. The engineered DNA polymerase and dNTPs with four fluorescent labels were added, and only one base was synthesised per cycle. The surface of the reaction plate was then scanned with a laser to read the nucleotide species polymerised in the first round of each template amplifications. The ‘fluorophore’ and ‘termination group;’ were then chemically cleaved to restore the 3’ end viscosity, and polymerisation to the second nucleotide proceeded. Finally, the fluorescence signal was counted for each round of amplification, and the sequence of the template DNA fragment was obtained.

#### Biological information analysis

PE reads obtained by Miseq sequencing were first spliced according to the overlap relationship, sequences were quality-controlled and filtered, and operational taxonomic unit (OTU) cluster analysis and species taxonomic analysis were performed after distinguishing samples [15].

Based on the results of OTU cluster analysis, a variety of diversity index and sequencing depth analyses were performed on the OUT dataset. Based on taxonomic information, statistical analysis of community structure was performed at each classification level. Based on the above analysis, a series of in-depth statistical and visual analyses of multi-sample community composition and phylogenetic information were performed using multivariate analysis and differential significance tests. Miseq sequencing analysis was commissioned by Shanghai Majorbio Bio-pharm Technology Co., Ltd (China).

### Quantitative analysis of TP, TN, TK and AK in soil

According to a previously described method [16], soil samples corresponding to plant samples (collected simultaneously) were thoroughly analysed after air-drying. The relative content of TP, TN, TK and AK was analysed using a TOC-VCPH organic carbon analyser, a Kjeldahl analyser, and a continuous flow analyser, respectively.

#### Quantitative analysis of TP

TP was determined by spectrophotometry at a wavelength of 400 to 490 nm. A 10.00 ml sample of digested solution of constant volume, filtration or clarification (V2 phosphorus 0.05–0.75 mg) was placed in a 50 ml volumetric flask, and 2 drops of dinitrophenol were added. The indicator was neutralised to a yellow colour by dropwise addition of 6 M NaOH, and 10.00 ml of ammonium vanadium molybdate reagent was added and the volume was adjusted with water (V3). After 15 min, the sample was measured at a wavelength of 440 mm using a cuvette of 1 cm diameter, and the instrument was zeroed with a blank test digestion liquid prepared as described above. Finally, a standard curve was used to establish a standard curve or linear regression equation, and the phosphorus concentration in the colour-developing solution was quantified from the calibration curve or regression equation.

#### Quantitative analysis of TN

In a sealed diffuser, soil samples were hydrolysed with a 1.8 M sodium hydroxide (NaOH) solution, and TN converted to an ammonia adduct under constant temperature conditions continuously diffused and was absorbed by boric acid (H3BO3). The content of soil hydrolysable nitrogen was calculated by titration with standard hydrochloric acid.

#### Quantitative analysis of TK

TK content in the test solution was determined by flame photometry. Cooking liquid (5.00–10.00 ml) in a 50 ml volumetric flask was diluted to the required volume using water, and the test solution was directly measured using a flame photometer and galvanometer. Finally, a standard curve or a linear regression equation was established and used to determine the concentration of TK in the test fluid.

#### Quantitative analysis of AK

Using a 1 M neutral NH4OAc solution as the extracting agent, NH^+^_4_ was exchanged with K^+^ on the surface of soil colloids, and the solution was missed with water-soluble K^+^. The AK content in the leachate was then directly determined using a flame photometer.

### Analysis of berberine content by HPLC

#### Instruments and reagents

An Agilent 1260 array detector was coupled to an Agilent SB C18 column (250 × 4.6 mm, 5 μm internal diameter, Agilent, USA) and an Agilent 1260 HPLC system. An appropriate amount of berberine reference substance (purity ≥99%, Beijing spectrum) was added to methanol to prepare a solution containing 20% per ml and used as a reference solution.

#### Sample preparation

Rhizome material was completely air-dried, made into a powder, and passed through a No. 4 sieve. Approximately 0.l g of powder was added to 50 ml methanol, densely packed, weighed, sonicated (power = 250 W, frequency = 40 kHz) for 1 h, cooled, weighed again, lost weight was replaced with methanol, and the sample was shaken, filtered, and the filtrate was used to prepare test solutions.

For chromatography, octadecylsilane-bonded silica gel (Agilent SB C18, 250 mm × 4.6 um, 5 μm internal diameter) served as a filler, 0.22 M acetonitrile and potassium dihydrogen phosphate solution (25:75) served as the mobile phase, and the detection wavelength was 265 nm. Sample and reference test solutions were 10 μl, and the berberine content was determined from the HPLC chromatograms.

### Statistical analysis

All data are expressed as mean ± standard deviation. Significant differences between groups were determined by t-tests. The analysis of similarities (ANOSIM) nonparametric test and Bray-Curtis algorithm were used to calculate distances between pairs of samples, and all distances were sorted in size order. R and R* values were calculated, and the probability that R* was greater than R gave the *p*-value (*0.01 <*p* <0.05, **0.001 <*p* <0.01, \*\*\**p* <0.001). The 16S rDNA gene sequencing data were analysed using the free online Majorbio I-Sanger Cloud Platform (www.i-sanger.com). The sequence data are available at the NIH Sequence Read ArchiveIn review (https://submit.ncbi.nlm.nih.gov/subs/bioproject/.) under the Bioproject accession number PRJNA556939.

## Results

### Quality metrics of pyrosequencing analysis

The average number of sequences cross all 30 samples was 903,568, the average number of nucleotides was 356,700,253 bp, and the average sequencing read length was 394.7685764 bp (Supplementary file 1). Based on the OTU results, we found that the number of OTUs in W and A groups was AR (1413) >WR (1102) >AS (1059) >WS (1003) >AL (901) >WL (816; Figure 2). The number of OTUs in cultivated leaf, stem and root samples was larger than in WT samples, and the number of OTUs was ordered roots > stems >leaves (Supplementary file 2).

**Figure 1.**
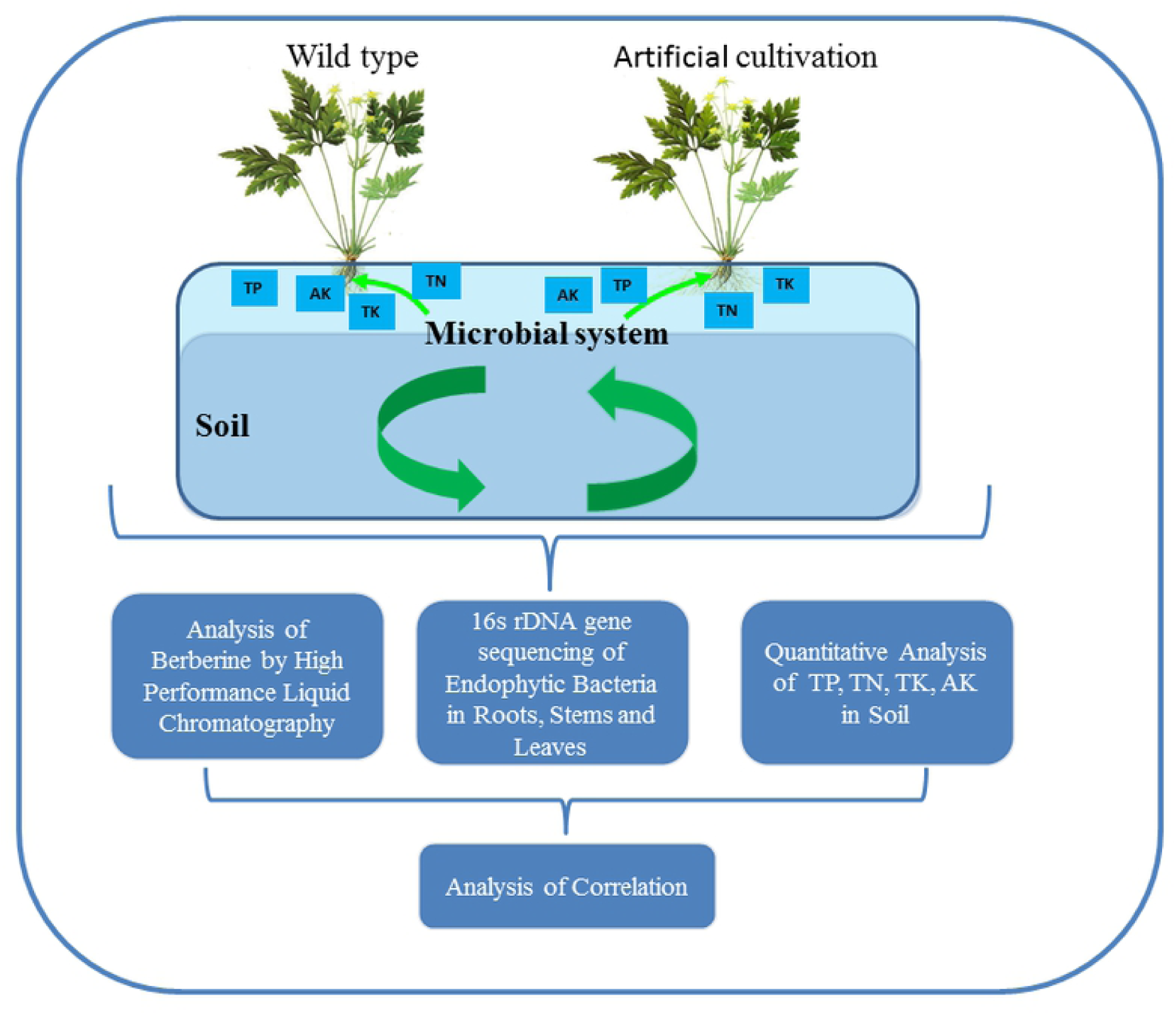
Flowchart of experimental processes.

**Figure 2.**
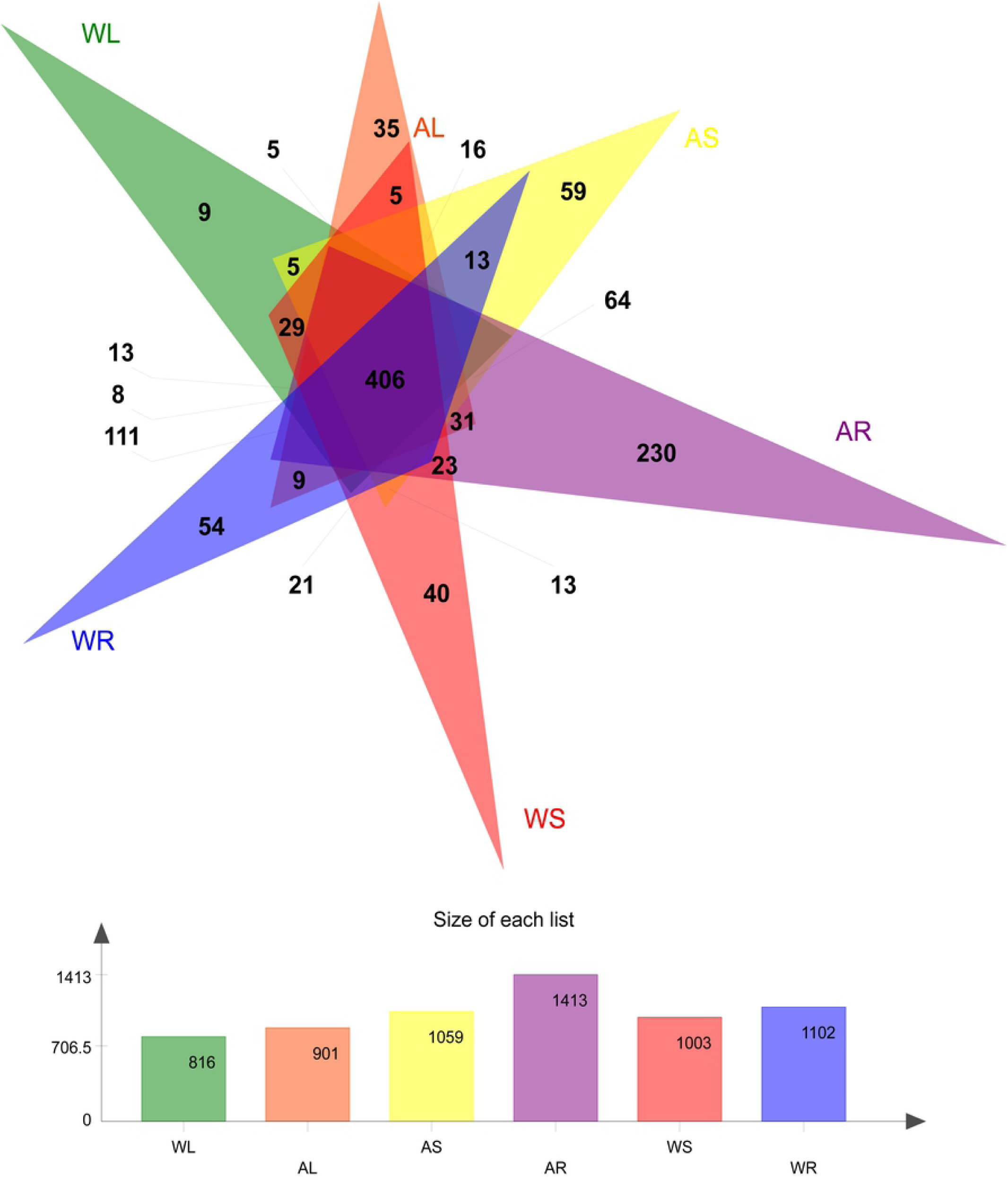
Venn diagram of identified OTUs. Different colours represent different groupings, overlapping parts represent species shared among multiple groups, no overlapping parts represent species unique to groupings, and numbers indicate the number of corresponding species.

### Alpha diversity

Based on the OTU results, we analysed Alpha diversity to investigate the diversity of microorganisms in the environment. This index can reflect the abundance and diversity of microbial communities using single sample diversity analysis, and it generates various statistical outputs. The analytical index estimates the species abundance and diversity of environmental communities, the Chao index reflects community richness, the Shannon index reveals community diversity, and the Shannoneven index reflects community evenness. The results revealed that the root endophytic Chao index of *Coptis teeta* Wall. was largest (*p* <0.01) and AR was greater than WR, indicating that the endophytic abundance of WT rhizoctonia solani was highest, and WT plants were superior to the artificial cultivar. Regarding the Shannon index, the value of WS for WT *Coptis teeta* Wall. was largest, while the value of AR in the artificial cultivar was largest, and the root endophytic Shannon index value of the artificial cultivar was greater than that of WT plants (*p* <0.01). These results showed that WT *Coptis teeta* Wall. exhibited high diversity for the WS group, while the artificial cultivar displayed high diversity for the AR group, and WT plants had higher root flora diversity than cultivated plants.

According to the Shannoneven index, the value for WS was largest for WT plants, while the Shannoneven value for the AR group was largest for the artificial cultivar, and the root endophytic Shannoneven value of the cultivar was greater than that of WT plants (*p* <0.01). These results indicate that the uniformity of the WS group in WT plants was higher, the uniformity of the flora in the AR group of artificially cultivated *Coptis teeta* Wall. was higher, and the uniformity of the root endophytic bacteria was higher for WT plants than for cultivated plants.

### Beta diversity

We selected the two evolutionary levels of OTU and genus to calculate beta diversity using principal component analysis (PCoA), hierarchical clustering (Hcluster) and ANOSIM. Additionally, PCoA analysis plots of 30 samples at the OTU level were generated based on the selected distance matrix (Figure 4a). Hcluster can clearly reveal the distances of sample branches, and a hierarchical clustering tree was generated based on the unweighted pair group method with arithmetic mean (UPGMA; Figure 4b). In addition, ANOSIM used the Bray-Curtis algorithm to calculate differences between pairs of samples, and to test whether differences between pairs of groups were significantly greater than intra-group differences (Table 1).

**Table 1.** Analysis of similarities (ANOSIM)

| Phylogenetic level<br>ANOSIM output | OTU |  | Genus |  |
| --- | --- | --- | --- | --- |
|  | R | P | R | P |
| WL VS AL | 0.396 | 0.024 | 0.42 | 0.024 |
| WS VS AS | 0.468 | 0.032 | 0.368 | 0.037 |
| WR VS AR | 0.58 | 0.012 | 0.524 | 0.01 |
| WR VS WL | 0.924 | 0.011 | 0.98 | 0.014 |
| WL VS WS | 0.648 | 0.009 | 0.624 | 0.008 |
| WR VS WS | 0.812 | 0.005 | 0.76 | 0.007 |
| AR VS AL | 0.992 | 0.011 | 0.996 | 0.007 |
| AR VS AS | 0.888 | 0.008 | 0.856 | 0.013 |
| AL VS AS | 0.8 | 0.01 | 0.82 | 0.012 |

The closer the R value is to 1, the greater the intra-group differences, and the smaller the R value, the less significant the differences between and within groups.

**Figure 3.**
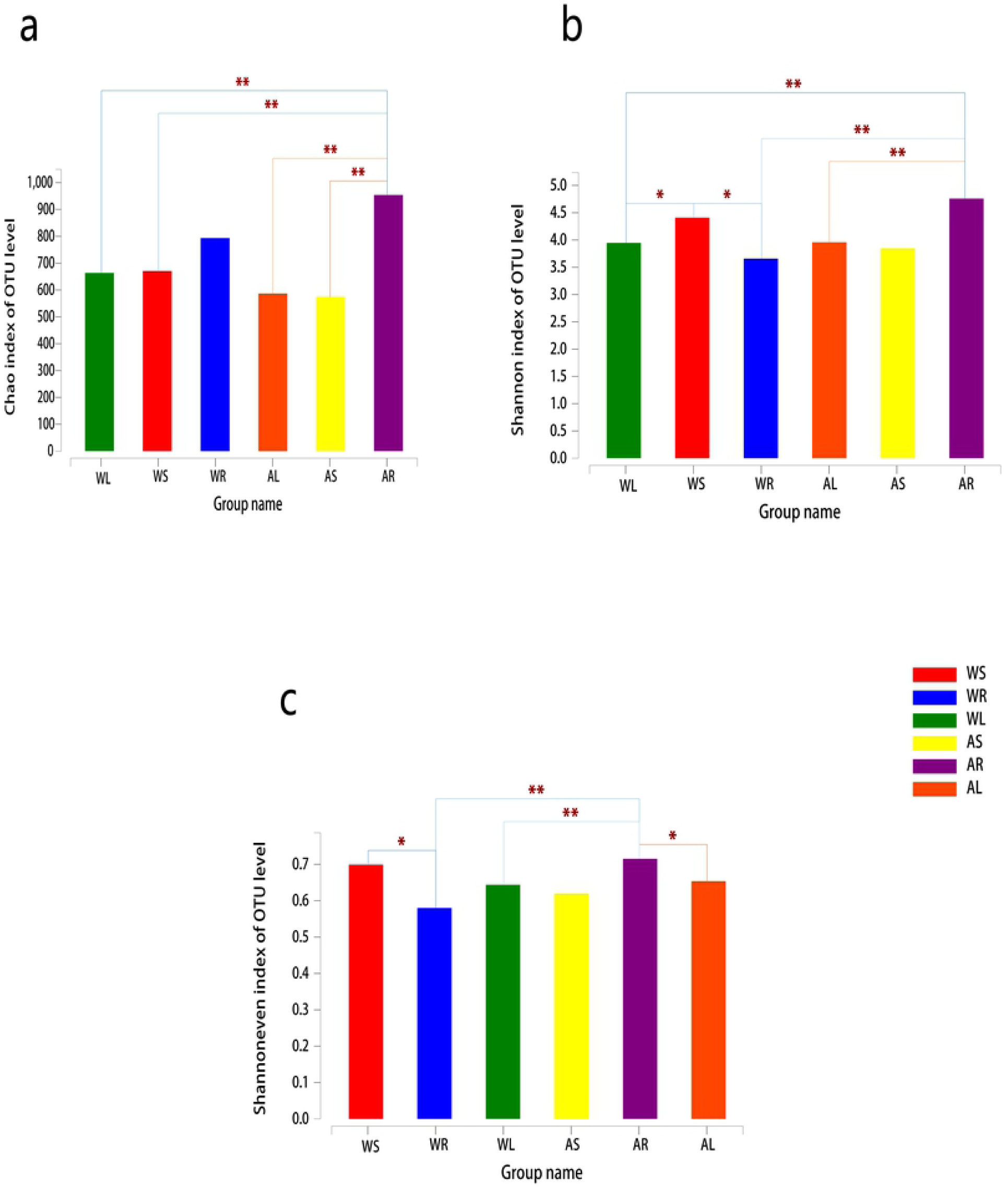
Alpha diversity estimates of bacterial communities. **a** OTU richness estimates (Chao index). **b** OTU diversity estimates (Shannon index). **c** OTU evenness estimates (Shannoneven index). The chart shows significant differences between the two groups of samples, and the two groups of markers with significant differences (*0.01 <*p* <0.05, **0.001 <*p* <0.01, \*\*\**p* <0.001). The abscissa is the group name and the ordinate is the exponential average of each group.

**Figure 4.**
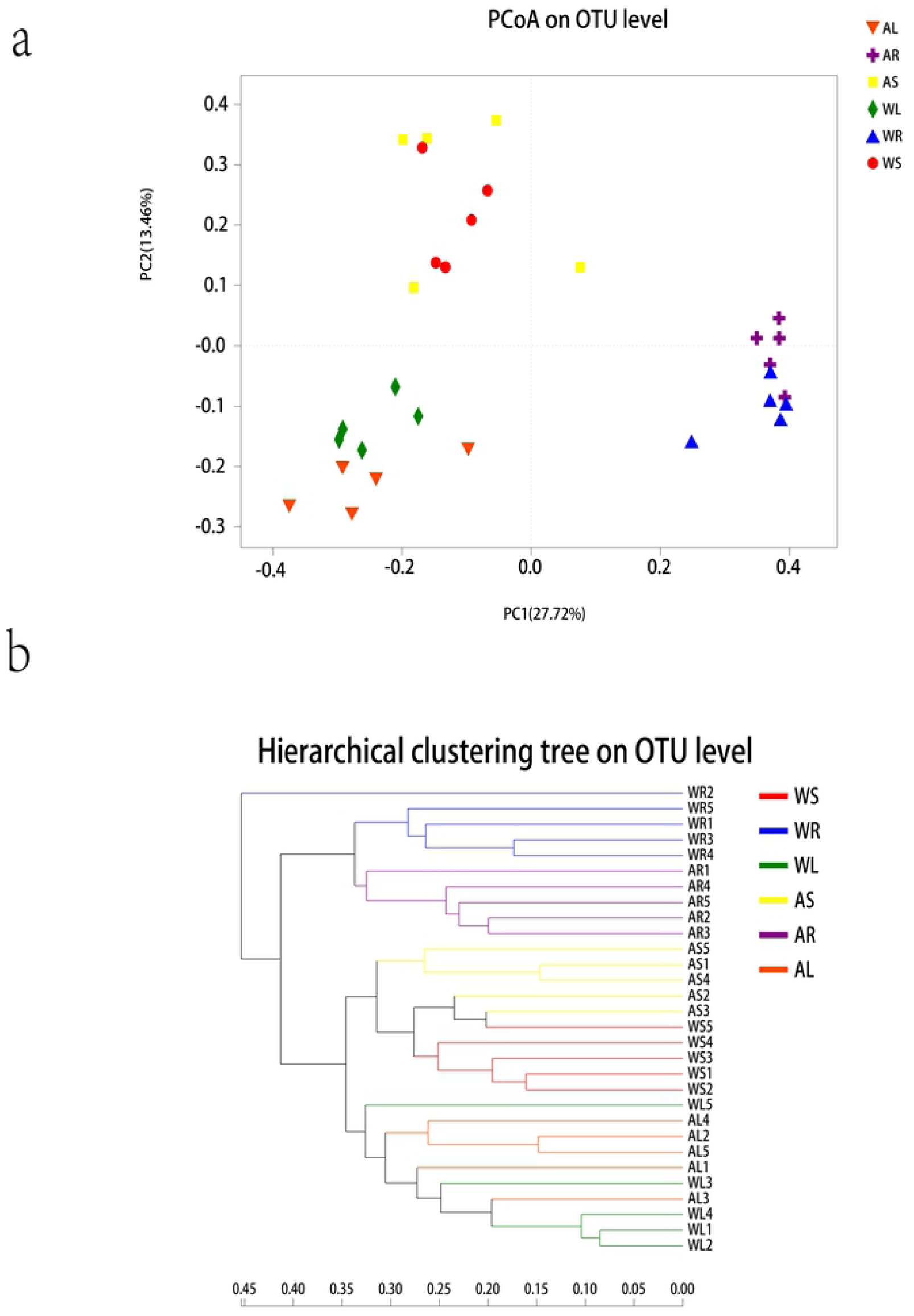
Beta diversity estimates of bacterial communities. **a** PCoA at the OTU level. Horizontal and vertical coordinates represent two selected principal coordinate components, and the percentage represents the contribution of the primary coordinate component to sample composition differences. Different coloured points and shapes represent samples from different groups, and the closer two sample points are, the more similar the composition of the two sample species. **b** Hierarchical clustering tree at the OTU level. The length of branches represents the distance between samples, and different groups are shown in different colours.

PC1 explained 27.72% of total variation at the OTU level, and PC2 accounted for 13.46% (Figure 4a). PCoA analysis based on unweighted UniFrac distances showed that endophytic bacteria in different parts of WT and cultivated plants differed considerably, and bacteria in roots, stems and leaves could be clearly distinguished (Figure 4a). Based on sample-level clustering, samples could be divided into six distinct groups according to bacterial OTUs (Figure 4b), further illustrating differences between WT and artificial cultivars (Figure 4b).

PCoA, Hcluster and ANOSIM results at the phylogenetic level (OTU and genus levels) further revealed differences in endophytes between different parts WT and cultivated plants (Table 1).

### Dominant members of the bacterial community in different plant tissues

We investigated the composition of bacteria in individual samples, and compared endophytic bacteria in the roots, stems and leaves of WT and cultivated plants using analysis of variance (ANOVA). According to compositional characteristics at the phylum level (Figure 5a), Actinobacteria (51.94%), Proteobacteria (43.85%), Firmicutes (2.27%) and Bacteroidetes (1.37%) were identified in WS, Proteobacteria (76.05%), Bacteroidetes (11.06%), Actinobacteria (11.08%) and Firmicutes (1.17%) were detected in WR, and Actinobacteria (70.12%), Proteobacteria (27.57%), Firmicutes (1.15%) and Bacteroidetes (0.57%) were present in WL. Meanwhile, Proteobacteria (62.39%), Actinobacteria (32.34%), Bacteroidetes (3.32%) and Firmicutes (0.85%) were identified in AS, Proteobacteria (70.01%), Actinobacteria (12.49%), Acidobacteria (3.4%), Firmicutes (0.80%) and Bacteroidetes (0.85%) were detected in AR, and Actinobacteria (62.93%), Proteobacteria (35.64%), Bacteroidetes (0.37%) and Firmicutes (0.46%) were present in AL. Thus, Phyla making the largest contributions were Proteobacteria, Actinobacteria and Bacteroidetes, while Firmicutes and Acidobacteria made minor contributions. Proteobacteria, Actinobacteria and Bacteroidetes varied widely between the six groups (*p*-values are shown in Figure 5b). Based on the results of compositional analysis at the genus level, *Mycobacterium, Salmonella, Nocardioides, Burkholderia-Paraburkholderia* and *Rhizobium* were the dominant genera (Figure 5c), and *Mycobacterium, Salmonella* and *Nocardioides* differed greatly among the six groups (*p*-values are shown in Figure 5d).

**Figure 5.**
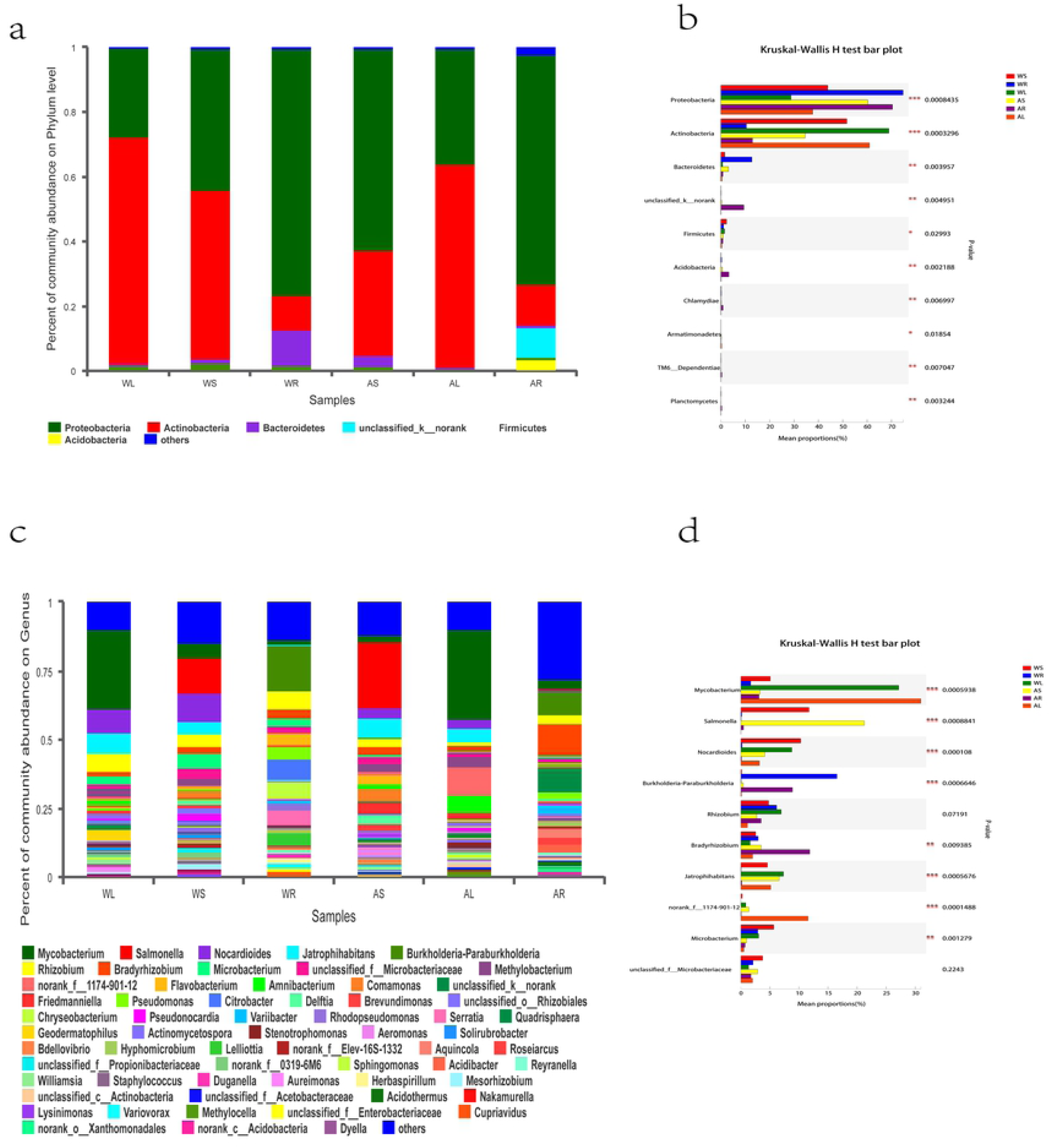
Distribution of OTUs at phylum and genus levels. **a** Percentage of community abundance at the phylum level. Different coloured columns represent different species, and the length of columns represents the proportion of species. **b** The top 10 dominant phyla. The vertical axis represents the species name at the phylum level, and the column length corresponding to the species represents the average relative abundance of the species in each sample group, with different colours indicating different groupings. The *p*-value is shown in the right (*0.01 <*p* ≤0.05, **0.001 <*p* ≤0.01, \*\*\**p* ≤0.001). **c** Percentage of community abundance at the genus level. Different coloured columns represent different species, and the length of columns represents the proportion of species. **d** The top 10 dominant genera. The vertical axis represents the species name at the genus level, and the column length corresponding to the species represents the average relative abundance of the species in each sample group, with different colours indicating different groupings. The *p-*value is shown on the right (*0.01 <*p* ≤0.05, **0.001 <*p* ≤0.01, \*\*\**p* ≤0.001).

### TK, TP, TN and AK in rhizosphere soil

We analysed the TK, TP, TN and AK content in WT and cultivated rhizosphere soil, and the results showed that the relative content of TK in WR was 7342 ± 1546.713 mg/kg, whilst in AR it was 5042 ± 1749.391 mg/kg. The relative content of TP in WR was 1098 ± 111.6692 mg/kg, compared with 972 ± 266.4958 mg/kg in AR. The relative content of TN in WR was 355.6 ± 34.55865 mg/kg, compared with 346.4 ± 37.38717 mg/kg in AR. The relative content of AK in WR was 1196.8 ± 284.2309 mg/kg, and the relative content in AR was 987.2 ± 131.648 mg/kg (Table 2).

**Table 2.** Analysis of TK, TP, TN and AK in rhizosphere soil (mean ± SD; mg/kg)

|  | WR | AR | N |
| --- | --- | --- | --- |
| TK | 7342 ± 1546.713 | 5042 ± 1749.391 | 5 |
| TP | 1098 ± 111.6692 | 972 ± 266.4958 | 5 |
| TN | 355.6 ± 34.55865 | 346.4 ± 37.38717 | 5 |
| AK | 1196.8 ± 284.2309 | 987.2 ± 131.648 | 5 |

TK, TP, TN and AK in rhizosphere soil.

### Berberine content in roots

Next, we analysed the berberine content in the roots of WT and cultivated *Coptis teeta* Wall. by HPLC, and berberine levels were significantly higher in WT roots (Figure 6).

**Figure 6.**
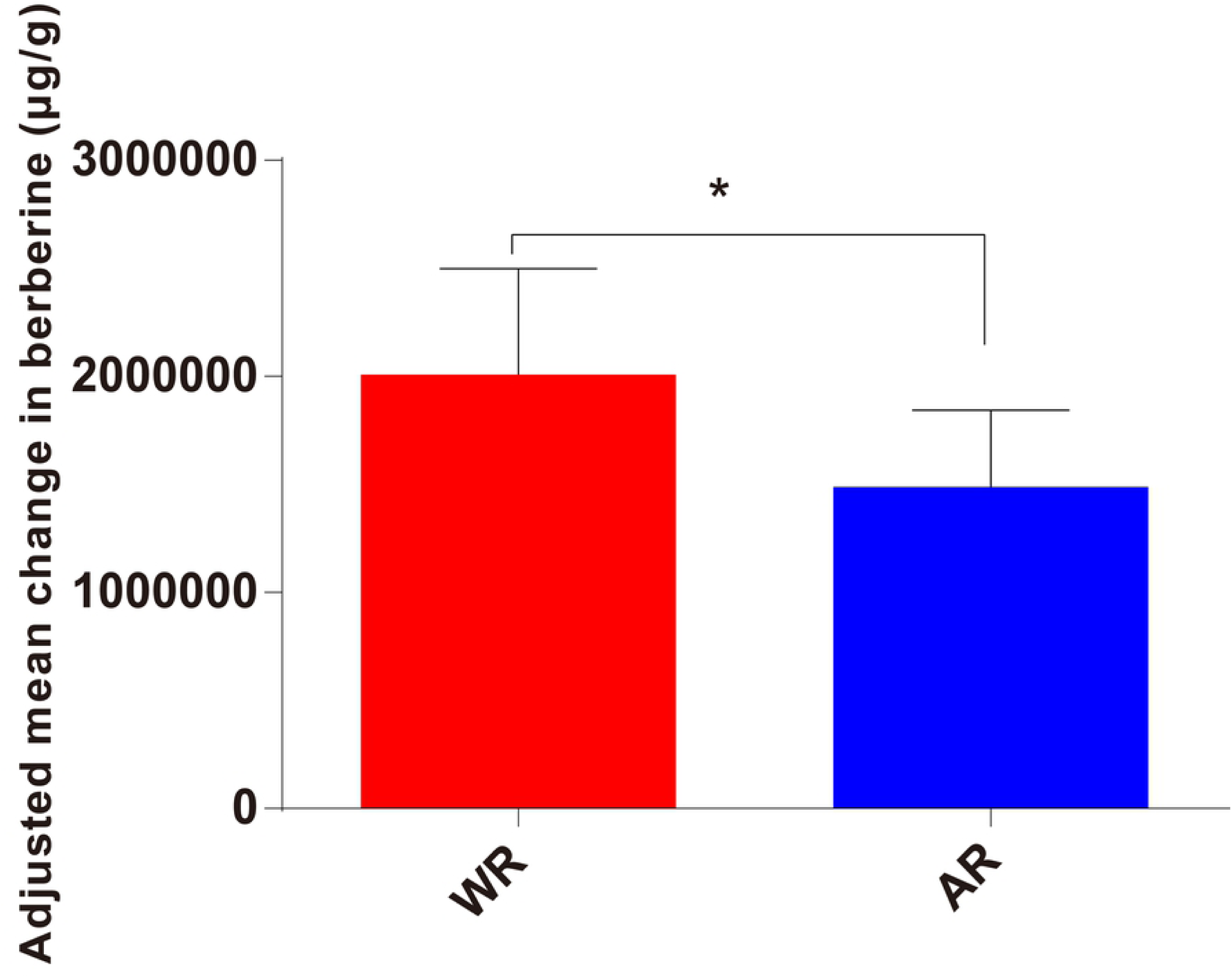
Content of berberine in roots. The chart shows significant differences between the two groups (*0.01 <*p* ⩽0.05).

### Correlation analysis

We used canonical correspondence analysis (CCA) to explore the correlation between endophytes in WT and cultivated *Coptis teeta* Wall. roots at the genus level, berberine content in roots, and TK, TP, TN and AK in rhizosphere soil. The results analysis showed that berberine had a significant positive correlation with TK, TP, TN and AK in rhizosphere soil (Figure 7a, b).

**Figure 7.**
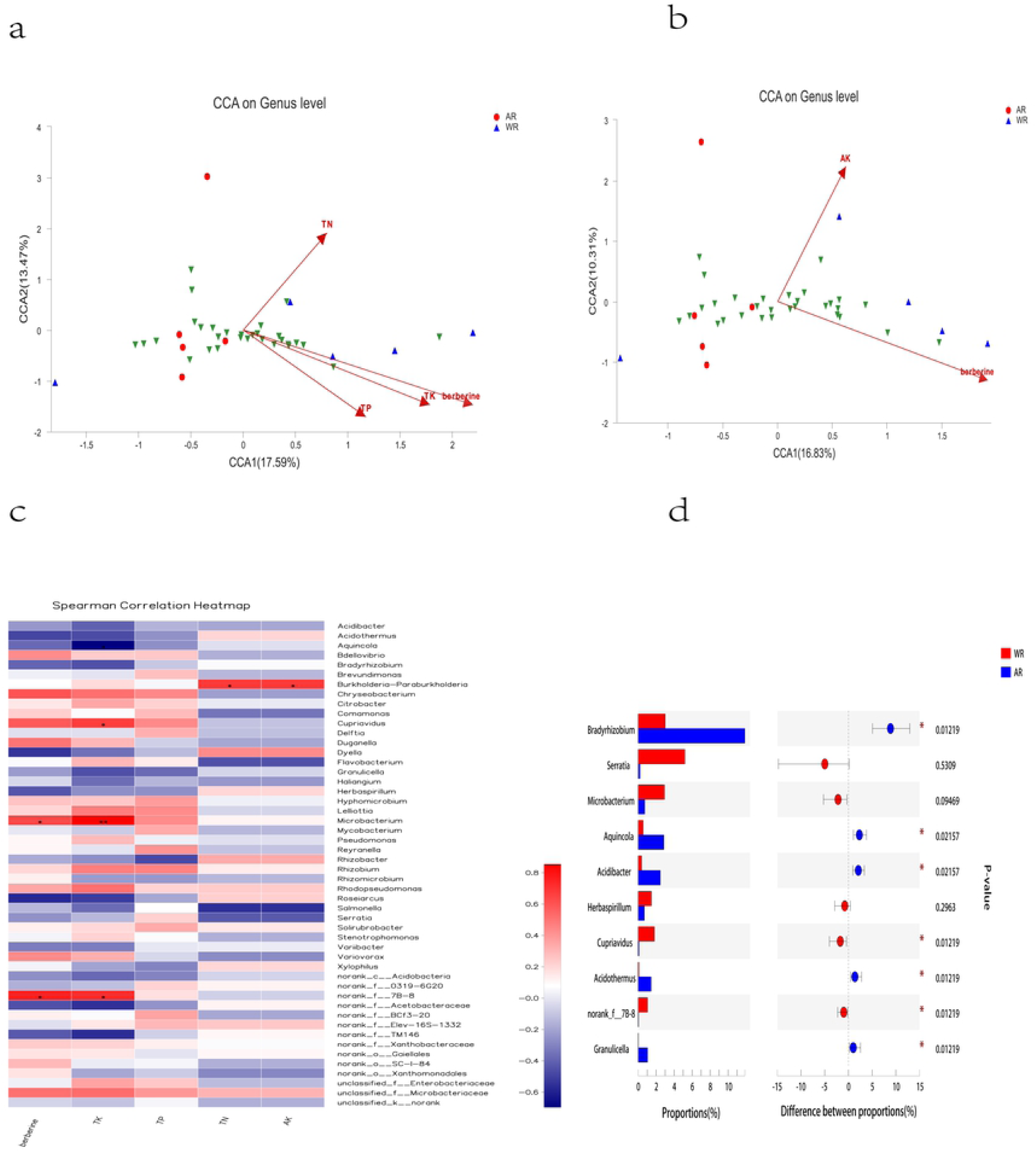
Environmental factor correlation analysis. **a** CCA at the genus level (1). b CCA at the genus level (2). Different coloured dots and shapes represent sample groups, green arrows represent species, red arrows represent quantitative environmental factors, and the length of environmental factor arrows represents the extent to which environmental factors affect the species data. The angles between environmental factor arrows represents the correlation (sharp angle = positive correlation, obtuse angle = negative correlation, right angle = no correlation). c Spearman correlation heatmap. The x- and Y-axes are environmental factors and species, respectively, and the R value and p-value were obtained by calculation. R values are shown in different colours, p-values <0.05 are labelled with an asterisk (*), and the legend on the right shows colour intervals for different R values (*0.01 <p ≤0.05, **0.001 <p ≤0.01, ***p ≤0.001). d The top 10 related genera. The vertical axis represents the species name at the genus level, and the column length corresponding to the species represents the average relative abundance of the species in each sample group, with different colours indicating different groupings. The p-value is shown on the right (*0.01 <p ≤0.05, **0.001 <p ≤0.01, ***p ≤0.001).

Finally, we used Spearman correlation analysis to assess the correlation between microbial classification and TK, TP, TN and AK in rhizosphere soil. We subjected 50 genera to in-depth analysis and found that berberine was positively correlated with *Microbacterium* and *norank_f_7B-8*. (*p* <0.05), TK was positively correlated with *Microbacterium* (*p* <0.01), and TN, AK and *g_Burkholderia-Paraburkholderia* were positively correlated (*p* <0.05; Figure 7c). Based on the correlation with berberine, combined with the characteristics of endophytic bacteria in WT and cultivated plants, 10 genera (*Bradyrhizobium, Serratia, Microbacterium, Aquincola, Acidibacter, Herbaspirillum, Cupriavidus, Acidothermus, norank_f_7B-8* and *Granulicella*) were identified that may be involved in the synthesis of berberine (Figure 7d).

## Discussion

The microbiomes of plants are receiving much attention form researchers, and knowledge is gradually being converted into field applications[17,18]. A better understanding of the relationships between plant microbial groups and soil environments, and the effects of probiotics on crop growth and development, could help us to improve the efficient absorption of nitrogen, phosphorus, iron and other elements by crops[19]. Plant-associated microbes can influence plant innate immune responses and resistance to a variety of environmental stresses[20], and knowledge could allow us to reduce the use of chemical fertilisers and pesticides whilst simultaneously greatly improving the yield and quality of plant products[21,22].The development of microbiology and related technologies will provide strong technical support for solving major problems in the sustainable development of agriculture, such as the abuse of pesticides and fertilisers, environmental pollution, and the rise of crop diseases. Therefore, it is urgent for us to investigate crop microbiomes to ensuring the safety of medicinal and food plants[23].

Previous studies have focused on indica and japonica rice varieties, and indica rice usually displays higher nitrogen use efficiency [24]. The diversity of indica rice root microflora is significantly higher than that of japonica rice, and characteristics of the root microflora may serve as biomarkers for distinguishing these two rice indica varieties. Interestingly, the roots of indica rice support more microbial species related to the nitrogen cycle than japonica rice, resulting in a more active nitrogen transformation environment, which may be one of the important reasons why the nitrogen use efficiency of indica rice is higher than that of japonica rice [24]. Wheat scab caused by *Fusarium graminearum* is an important crop fungal disease that reduces wheat yield over large areas. In addition, mycotoxins secreted by *F. graminearum* can accumulate in infected wheat grains, threatening human and animal health, and posing serious food safety problems. *F. graminearum* mainly infects the spikes of wheat during the flowering stage, and initiation of infection is dependent on the identification of hosts [25]. Sesame is a traditional medicinal plant and popular green leafy vegetable. Sesame is colonised by a diverse, habitat-specific microbiome, with an indigenous interleaf bacterial community dominated by Enterobacteriaceae, and a variety of antibiotic resistance traits have been observed [26].

In our present study, based on overlap between PE reads, pairwise reads were spliced, merged, filtered, sequenced using barcodes, and the sequence direction was corrected to generate optimised data. The quality metrics of pyrosequencing analysis indicated uniform, high-quality sequencing suitable for subsequent analysis [27].

The number of OTUs identified in cultivated leaf, stem and root samples was larger than in WT samples, and the number of OTUs both WT and cultivated plants was ordered roots >stems >leafs. Soils are among the most abundant microbial ecosystems on earth. Endophytic bacteria can enhance the decomposition of nitrogen, phosphorus, potassium and other nutrient elements in soil [28], and in general soil closer to roots yields a greater number of bacterial OTUs. Some artificial cultivars can control infection, adjust nutritional structure, and may possess superior disease resistance [29], consistent with a larger number of OTUs in cultivated vs. WT plants.

According to the results of alpha diversity (Chao, Shannon and Shannoneven indices), we found that the abundance, diversity and evenness of endophytic bacteria ere greater in WT than in artificial cultivated plants. This may be related to the fact that *Coptis teeta* Wall. is a shade-tolerant plant that grows at high altitude, making it suitable for cold mountainous and rainy areas, and it has distinct environmental requirements. Therefore, the diversity, abundance and evenness of endophytic bacteria associated with WT *Coptis teeta* Wall. were greater than those of cultivated plants.

Differences may be explained from another aspect. The results of beta diversity analysis focusing on PCoA, hierarchical clustering (Hcluster) and ANOSIM revealed significant differences in endophytic bacteria between WT and cultivated plants, and between roots, stems and leaves. The plant root system is the interface between these multicellular eukaryotes and soil [30,31]. The Interleaf and associated microorganisms constitute a large and dynamic ecosystem that is important at a global scale [32].

We found that the dominant phyla were Proteobacteria, Actinobacteria and Bacteroidetes, and the most abundant genera were *Mycobacterium, Salmonella, Nocardioides, Burkholderia-Paraburkholderia* and *Rhizobium*. These results revealed details of the composition structure and characteristics of this plant species. Compositional differences at phyla and genera levels could potentially be used to distinguish between WT and artificial cultivars, and may be related to the absorption of nutrients from soil and the biosynthesis of pharmaceutical compounds.

We measured the content of TK, TP, TN and AK in rhizosphere soil, and all were more abundant in rhizosphere soil of WT plants. Soil pH is an important factor determining the diversity and structure of nitrogen-fixing bacterial communities [33]. Endophytic bacteria can enhance the decomposition of nitrogen, phosphorus, potassium and other nutrient elements by host plants [34]. Nitrogen is an indispensable component of all organisms, and it is also necessary for the synthesis of key cellular compounds such as proteins and nucleic acids [35]. Nitrogen in the atmosphere is the largest stock of freely available nitrogen, but few bacteria can fix this inert gas [33]. Therefore, these characteristics may be related to the synthesis of medicinal components.

There are many medicinal components present in *Coptis teeta* Wall. [36]. Herein, we analysed the content of berberine in the roots of WT and artificially cultivated plants by HPLC, and quantities in WT roots were higher than in artificially cultivated plant roots. WT *Coptis teeta* Wall. can freely absorb nutrients from the soil and take full advantage of the natural environment, which may result in superior biosynthesis of compounds such as berberine. Therefore, endophytic bacteria might be related to the synthesis of berberine and TK, TP, TN and AK levels in rhizosphere soil.

Our results revealed that berberine was positively correlated with TK, TP, TN and AK levels in rhizosphere soil of *Coptis teeta* Wall.. Berberine was positively correlated with *Microbacterium* and *norank_f__7B-8*, TK was positively correlated with *Microbacterium*, and TN, AK and *g__Burkholderia-Paraburkholderia* were positively correlated (Figure 7c). These findings indicate that endophytic bacteria may be related to the soil environment. Under certain circumstances, soil nutrients can affect the composition and structure of endophytes [37].

Furthermore, based on the correlation with berberine and the characteristics of endophytic bacteria in WT and artificially cultivated *Coptis teeta* Wall., 10 genera (*Bradyrhizobium, Serratia, Microbacterium, Aquincola, Acidibacter, Herbaspirillum, Cupriavidus, Acidothermus, norank_f__7B-8* and *Granulicella*) were identified that may be involved in the synthesis of berberine (Figure 7d). Some of these genera may be directly linked to berberine synthesis, and some may interfere with berberine synthesis since antagonism between bacteria the balance between positive and negative factors affecting berberine synthesis. For example, *Bradyrhizobium* species are believed to utilise sugars and organic acids, and studies have found that this genus is resistant to antibiotics.[37] *Serratia* are found in soil, water, and on the surfaces of plants, and some members of this genus are conditional pathogens of humans [38]. *Microbacterium* strain EC8 can enhance root and shoot biomass in lettuce and tomato [39], and many studies have shown that members of this genus can produce the plant growth hormone indoleacetic acid and solubilise phosphate [40]. Additionally, *Microbacterium* isolated from soil suppressed Rhizoctonia root rot in wheat, and enhanced the growth of wheat seedlings and reduced root infections [41]. In the present work, *Microbacterium* and *norank_f__7B-8* were strongly correlated with the synthesis of berberine, and may explain why WT *Coptis teeta* Wall. produced more berberine than artificially cultivated plants [42]. However, further verification is needed to explore the production of berberine in this plant species.

Herein, we identified endophytic bacteria related to berberine in the Chinese medicinal plant *Coptis teeta* Wall. using 16S rDNA sequencing and metabonomics. The microbial composition of WT and cultivated plants differed, and microorganisms in the root, stem and leaf varied. Furthermore, our results demonstrated a clear correlation between endophytic bacteria and berberine production. The findings provide a basis for detailed studies on the precise endophytic bacteria essential for berberine production, and may improve berberine production, as well as food and medicine safety.

## Supporting Information

Supplementary file 1: Sequence information table; Supplementary file 2: OTU information table.

## Data Availability

The data used to support this study can be made freely available.

## Author’s Contribution

Li-guo Chen, Jia-li Yuan, and Shou-zheng Tian participated in its design and searched databases and helped to draft the manuscript. Tian-hao Liu, Xiao-mei Zhang carried out the statistical analysis of data.

